# Anti-microbial activity of potential probiotic lactic acid bacteria against Methicillin-Resistant Staphylococcus Aureus (MRSA)

**DOI:** 10.1101/2020.03.08.982512

**Authors:** Alazar Essayas, Sujata Pandit, Pankaj Taneja

## Abstract

Lactic acid bacteria (LAB)are the essential ingredients in probiotic foods, intestinal microflora, and dairy products able to cope up and exist in diverse environmental ranges. Samples were collected using sterile test tubes and transported to a laboratory in the icebox for further biochemical characterization. Gram test and catalase activity were examined after microscopically distinct colonies were sub-cultured to pure colonies based on standard gram and catalase test protocols. Subsequently, these bacteria were characterized for their ability to grow at various salt concentrations (5%,10%, and15%) and temperature gradients (15°C, 30°C, 45°C). Acid-tolerance was analyzed by growing the colonies in MRS broth adjusted to acidic pH (pH 3) and pH 7.2 (control). Bile tolerance of LAB isolates was assayed by growing in 0.3% bile-supplemented MRS agar. Bile salt hydrolase (BSH) activity was studied by growing 10 μl of the prepared overnight culture on BSH screening media containing MRS agar plate supplemented with bile salts. The LAB isolates were checked for antimicrobial activity by agar well diffusion assay. All isolates found gram-positive, catalase-negative and non-motile, convex elevation and entire margin. All LAB isolates were able to grow at 5-10% Nacl concentration, whereas moderately grow at 10% concentration but rarely grow at 15% Nacl concentration. BCM2, BBM3 and BGM1 record the highest acidic resistance viability percentage 94.9%, 92.7%, and 91.8% respectively. BCM3 has the lowest acidic resistance with a viable percentage of 87.4%. BBM1 records the highest bile tolerance activity whereas BCM2 has the lowest bile tolerance. All isolates were found BSH positive. The study reveals LAB isolates showed a putative probiotic potential.

**Article highlights:** LAB are the main ingredients of probiotic products commercially available in the market nowadays. To effectively functioning the host gastrointestinal tract probiotics, need to have certain criteria like acid and bile tolerance this study reveals

- Acid tolerance activity of LAB isolated from bovine milk
- Bile tolerance activity of LAB isolated from bovine milk
- Bile salt hydrolase (BSH) activity of LAB isolates
- Antimicrobial activity of LAB against MRSA

## 1.0 Introduction

Lactic acid bacteria(LAB) are gram-positive, catalase-negative, either rod or cocci-shaped, strictly fermentative and non-sporulating microbes with lactic acid as the main end product of carbohydrate fermentation [1]. It plays significant role in agricultural, food and clinical industries. Bacteriologists classify them under four genera, *Lactobacillus, Leuconostoc, Pediococcus, and Streptococcus*. Recent taxonomic revisions have proposed the following genera under the LAB category*, Aerococcus, Alloiococcus, Carnobacterium, Dolosigranulum, Enterococcus, Globicatella, Lactococcus, Oenococcus, Tetragenococcus, Vagococcus, and Weissella*. LAB’s application is mainly related to their safe metabolic activity in food products that produce organic acids and other important metabolites that help to increase the shelf life of food. Their abundance in various ecological niches and preservative impact on food makes them universally accepted as GRAS (Generally Recognized as Safe) for human consumption [2]. LAB naturally found in soil, plants, and predominantly in the intestines of herbivorous animals. 100 trillion micro-organisms colonize the intestines of mammals as micro-flora, which is important for the health of hosts [3]. Intestine naturally enclose bacteria, which are often described to as our ‘normal flora’ which comprises more than 400 bacterial species with many desirable qualities [4]. The intestinal microbiota creates a barrier for pathogens. *Lactobacillus and Bifidobacterium* species origin from human intestine produce antimicrobial molecules that are active *in-vitro* and *in-vivo* against enteric pathogenic microorganisms causing diarrhea. Nowadays, the modern world is skeptical about chemical preservatives and processed foods, but the public has trust in the LAB as a safe way for food preservation and promoting health [5]. Most fermented foods such as cheese, yogurt, pickled vegetables mainly produced through fermentation by LAB. In earlier days, food was fermented by providing the optimum environment for microbes naturally present in the food, whereas these days, starter culture products are supplemented by the introduction of selected novel strains with well-defined properties[2].

Probiotics are “live microorganisms which confer a health benefit on the host, when administered in adequate amounts”. Probiotic research got attention as scientific evidence keeps growing about its properties, function and beneficial effects of probiotic bacterial species in human livelihood. Probiotics consumption began since the history of man using fermented foods in ancient Greeks and Romans. High demand for probiotics, functional foods and dietary supplements due to rise of public awareness about intestinal health. It is now well understood that some of the infections and disorders such as irritable bowel syndrome, inflammatory bowel disease, and antibiotic-induced diarrhea could be treated with the use of probiotics [6]. Their benefits include alleviation of lactose intolerance, enhancing nutrients bioavailability, and prevention or reduction of the prevalence of allergies in individuals with high susceptibility[7]. Probiotics need to have certain potential, such as the ability to tolerate GIT passage, adherence to mucus and human epithelial cells, competing for space against pathogenic microbes through aggregation and inhibition of enteric pathogens [8]. LAB and *bifidobacterium* are commonly used in commercially available probiotic products. Traditionally, fermented foods serve as good sources of new potential sources of LAB. Therefore, the use of strains from these sources would have great importance to consumers suffering from lactose intolerance. To function effectively, A probiotic strain need to withstand and survive gut passage of humans to enter the site in a viable state and should be enough in their population (normally 10^7^ cells per ml). The bacteria should, therefore, exhibit a high tolerance to low (acidic) pH and good resistance to high concentrations of bile salt to survive and colonize the intestine[9]. Bile is also another factor that affects the viability of bacterial cell and it’s interaction with the environment since lipids and fatty acid components are likely to be degraded by bile salt [10]. Bile is a yellow-green solution comprising bile acids, cholesterol, phospholipids and biliverdin pigment. It is produced in the pericentral liver hepatocytes, inter-digestively accumulated and processed in the gallbladder, and released into the duodenum after food ingestion. Bile acts as a biological detergent that emulsifies and solubilizes lipid and therefore plays key role in fat digestion. Bile also provides strong antimicrobial activity, mainly by dissociation of membrane lipids. Bile acids synthesized in the liver from cholesterol, cholic and chenodeoxycholic acid, de novo [11]. Gastrointestinal tract was considered the ideal origin for probiotic strains since it was recommended to use host origin bacteria to successfully colonize the human/animal GIT tract [12]. LAB produce bacteriocins, namely, polypeptides synthesized ribosomally by bacteria that can have a bactericidal or bacteriostatic effect on other competitive bacteria [13]. Bacteriocins generally lead to cell death due to inhibiting biosynthesis of a cell wall or by damaging the membrane by creating pores [14]. Therefore, in food fermentations, bacteriocins are important in preventing food spoilage due to the inhibition of food pathogens. Well studied bacteriocin is nisin, which is widely used in food preservation and often used as additive especially in processed cheese, milk products, and canned foods[15]. Probiotic dietary products which shared (60-70%) of the global functional food market, are widely accepted as a major functional food market by customers around the world[16]. Many probiotics products introduced to the market without sufficient analysis of the probiotic effects of the strains, because these products are species specific and the product’s performance is closely linked to the product’s specific strains. Appropriate studies on the properties of LAB should be done before market release [6]. Commonly, the initial requirement of LAB with probiotic potential focuses on their resistance to acid and bile in the gastrointestinal tract (GIT) and their potential to colonize the host [17]. The objective of this study is evaluating the acid and bile tolerance activity of LAB isolated from bovine milk samples.

## 2.0 Materials and Methods

### 2.1 Sampling

A total of 30 Dairy samples taken from nearby dairy farms (Diary Treatment Farm and Bahubali Milk Farm) in Greater Noida, Delhi, NCR, India. Bovine milk samples from cow, buffalo and goat were collected on December 18, 2018, during the morning milking session at 6 AM. Samples collected using a sterile tube and brought to the laboratory in an icebox and kept in a refrigerator until examination. Samples were coded as BCM1, BCM2, BCM3, BBM1, BBM2, BBM3, BGM1. BCM stands for milk from a bovine cow, BBM refers to milk from buffalo’s and BGM stands for bovine goat’s milk.

### 2.2 Enumeration and isolation of lactic acid bacteria

In 9 ml of 0.9 percent sterilized saltwater, 1 ml of each milk sample was homogenized. For each sample, a 10-fold serial dilution (10^−1^ to 10^−6^) prepared using 1 ml of homogenate. The dilution of 100 microliters was distributed over low pH DeMan Rogosa Sharp (MRS) agar and incubated at 37 °C for 48 hours. Colonies with typical LAB characteristics selected randomly and sub-cultured to pure colonies after primary microscopic and macroscopic examinations. Colonies with LAB characteristics were chosen for further physiological and biochemical studies.

### 2.2 Identification of LAB

#### 2.2.1 Gram staining

A modified gram stain protocol by Brucker [18] was conducted. Microscopic slides were cleaned with 95% ethanol and a drop of distilled water poured in the middle of the slide and bacterial colony smeared with flame sterilized inoculating loop.

After heat-fixing samples by hovering over the flame, gram staining was performed using crystal violet, iodine, 95% ethanol and safranine respectively. Colonies microscopically examined using 100 × oil immersion microscope. Purple bacterial samples were considered gram-positive and the pink-colored were identified as gram-negative bacteria.

#### 2.2.2 Catalase test

A modified methodology by Taylor [19] used for catalase test procedure. The clean microscope slide was prepared and distill water poured in the middle of a clean slide. The Bacterial colony picked using a flame-sterilized inoculating loop and colonies smeared on the water drop. 3% H_2_0_2_ was dropped on the sample to observe bubble formation or not due to catalysis of H_2_0_2_.

#### 2.2.3 Motility test

A semi-solid agar medium prepared in a test tube and isolated colonies inoculated using straight wire and stabbed down the center of the tube to the half depth of the medium. Samples incubated at 37°C for 48 hours. Cell motility was assayed by observing turbidity formation[20].

#### 2.2.4 Sodium chloride tolerance test

Sodium chloride (NaCl) tolerance was studied by growing LAB isolates in MRS broth adjusted with various NaCl concentration. NaCl in 1% (0.4 grams of sodium chloride in 40 ml of broth), 5% (2 grams of sodium chloride in 40 ml of broth) and 10% (4 grams of sodium chloride in 40 ml of broth) were prepared.50 microliters of LAB isolates inoculated to each salt concentration in 5 ml of the MRS-NaCl-containing tube and incubated for 48 hours. Cell viability was studied by subculturing the isolates to agar medium for colony formation

#### 2.2.5 Temperature tolerance test

LAB isolates growth pattern to different temperature ranges (10°C, 15°C and 40°C) was studied to analyze temperature tolerance.

#### 2.2.6 Carbohydrate fermentation test

Isolated LAB was tested for fermentation of sucrose, lactose, mannitol, dextrose, and maltose as per standard procedure.

### 2.3 Acid tolerance activity

A modified methodology by Liong and shah [21] used to study acid tolerance activity of LAB isolates. LAB isolates inoculated to normal saline water adjusted to pH3.0and pH 7.2(control). The pH of the broth adjusted using 1N HCL and distributed to the autoclaved test tubes. 1 percent (50 micro-liters) of cells inoculated in to the 5 ml broth and incubated at 37°C for 24 hours. Spectrophotometer was used to quantify the optical density of the sample. Tenfold serial dilution was prepared up to 10^−6^ to each bacterial strain after incubation using phosphate buffer saline (PBS). 100 micro-liters from each dilution was spread on MRS agar and incubated for 48 hours at 37°C.The colony count was enumerated as CFU / ml. Acid tolerance was calculated by comparing viable cell count at 0 hr. and 24 hrs. experiment was done in triplicate.

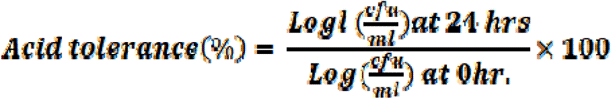

### 2.4 Bile tolerance activity

Bile tolerance activity studied using a modified methodology of Liong and Shah[21] with minor modifications. 1% of isolate were inoculated into 0.3% bile supplemented MRS broth and incubated at 37° C for 24 hours, simulating bile salt concentration of 0.3% in the GI tract. The optical density of each isolate was recorded at time 0 (t=0 hr.) and at time 24 (t=24hrs). Bile tolerance activity was calculated as

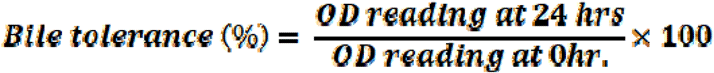

### 2.5 Bile salt hydrolase activity

Bile salt hydrolase activity was conducted as described by [22]. Bile salt hydrolase (BSH) activity was screened by growing 10 μl of the prepared overnight culture on MRS agar plate supplemented with 0.37 g/l CaCl_2_ with 5g/l sodium salt of taurodeoxycholic acid. Plates incubated at 37°C for 3 days. White precipitation around colonies considered as positive result. Experiment was done in triplicate.

### 2.6 Anti-microbial activity

Antimicrobial activity of potential probiotic LAB isolates was assed using agar well diffusion assay. MRSA concentration was adjusted to 0.5 MacFarland standard. 100 μl of S. aureus spread with a sterilized cotton swab on Muller Hinton agar plates and 100 μl of isolates were filled to a uniformly punched well. Inoculated plates incubated at 37 ºC for 48 hours and zones of inhibition measured (in mm) using a ruler. Experiment was in triplicate

### 2.6 Storage of LAB in a glycerol stock

100 micro-liters of 0.9% of saline water was used to a sterilize eppendorf tube and a colony of bacteria from overnight fresh cultures using a flame sterilized loop into 0.9% saline water and 500 micro-liters of 20% glycerol solution added to eppendorf tube and stored at−20 °C Freezer in Sharda University, Department of biotechnology, laboratory.

### 2.7 Statistical analysis

All statistical analyses were conducted using SPSS software version 20.0. All the experiments were conducted in triplicate and data was analyzed and compared statistically using ANOVA at a 95% level of significance. A probability value at p ≤ 0.05 was considered statistically significant. Data presented as mean values ± standard deviation calculated from triplicate experiments.

## 3.0 Result

A total of 7 LAB isolates were isolated and characterized from bovine milk samples. Table 1A shows the morphological, physiological and biochemical characteristics of LAB isolates. All isolates were found gram-positive, cata-lase-negative and non-motile. From isolated probiotic LAB strains BCM1, BCM3, BBM3, and BGM1 have cocci creamy white colonies on the MRS agar plate. BBM1 was the only tetrad structured LAB isolates from seven. All isolates were found convex in their elevation and entire margins BCM1, BBM1, BBM2, BBM3, and BGM1 have a smooth surface under microscopic observation whereas others have rough surface colonies. All the isolates were able to grow at 15°C, 30°C and 45°C.all isolates were positive for carbohydrate fermentation test (Table 1B) In this study, all the isolated probiotic isolates were able to grow at a concentration of 5-10 % NaCl, whereas fairly grow at 10% salt concentration, but rarely grow at 15% concentration (Table 2)

BCM2, BBM3 and BGM1 was the highest acidic resistance viability percentage 94.9%, 92.7%, and 91.8% respectively, whereas BCM3 has the lowest acidic resistance viability percentage of 87.4% (Figure 1 and Table 3) As shown in Figure 2, BCM1 and BBM2 record an increase in their OD reading after a 24-hour incubation in 0.3% bile supplemented MRS broth while the rest show a decrease in their OD reading and still viable using spectrophotometer. BBM3 records the highest viability percentage with 88.2% and BCM2 has the lowest viability percentage with value 33.3%.

**Fig. 1.**
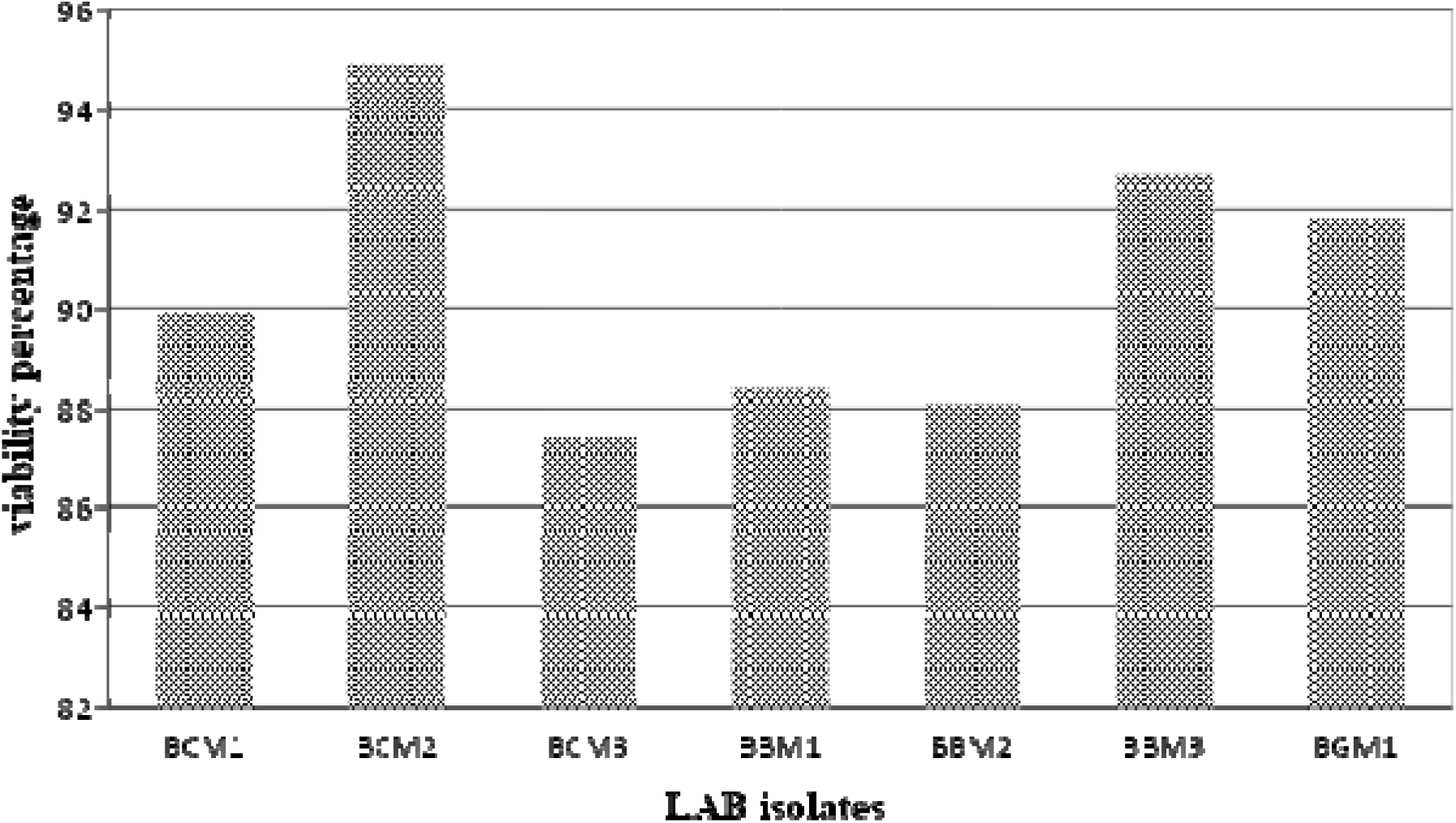
The viability percentage of LAB isolates to acidic pH (Ph=3) at 37° C when compared to pH=7. Data are expressed as mean ± SD (n = 3).

**Fig. 2.**
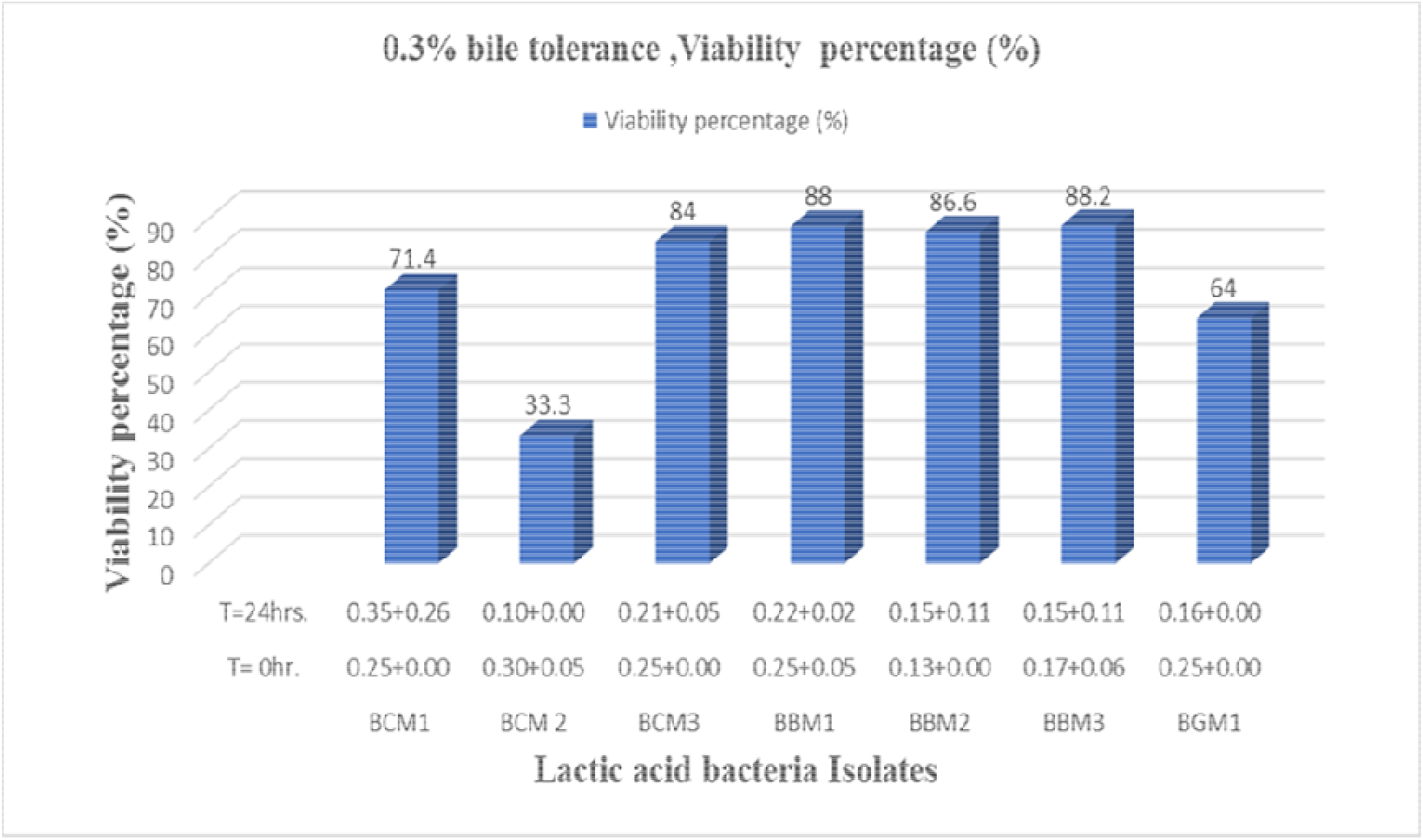
Bile tolerance percentage when compared at time t=0 hr. And time t=24hrs. Data expressed as mean ± SD (n = 3).

All the 7 isolates found BSH positive, they have the ability to hydrolyze bile salt after growth in Bile salt supplemented MRS media and exhibiting white opaque colonies (Table 4) All seven isolates of Lactobacillus had antimicrobial effects against MRSA, but the degree of antagonism among the isolates of Lactobacillus varied. BGM1 has the highest antagonistic activity with 29 mm zone of inhibition, whereas BCM2 has the lowest antibacterial activity (11 mm) against methicillin-resistant staphylococcus aerues. BBM1 is the second highest antimicrobial activity showed (Figure 3).

**Fig 3.**
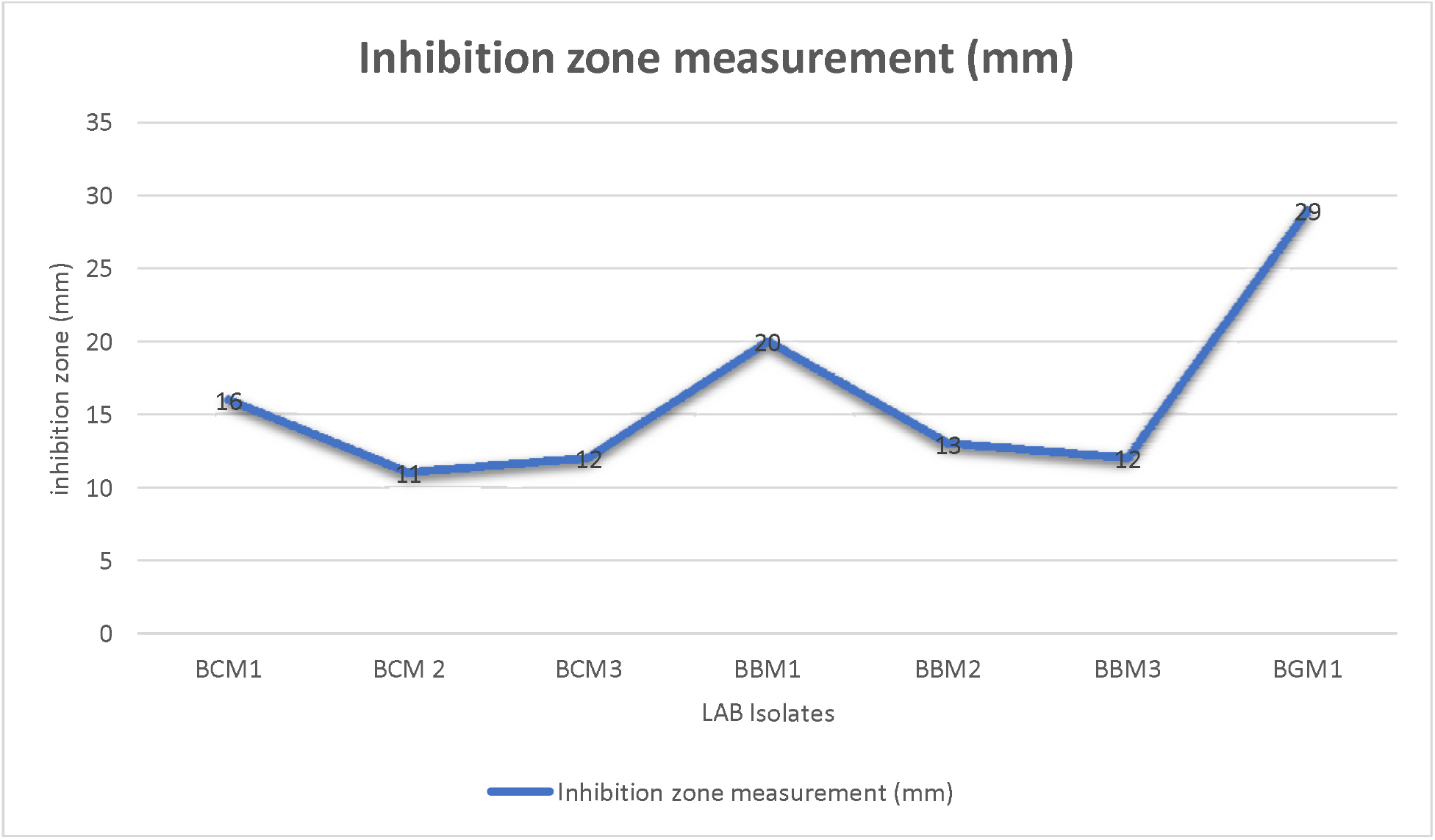
Antimicrobial activities of neutral pH CFS of LAB isolate against S. aureus, Data is expressed as mean ± SD (n = 3).

## 4.0 Discussion

LAB is a group of bacteria shaped of rod or coccus, gram-positive, catalase-negative, non-motile, homofermentative or heterofermentative, and able to grow in low acid condition [1], [23]. Morphological, physiological and biochemical characteristics of this study is aligned with LAB properties. Carbohydrate fermentation showed positive for all isolates. (Table 1 B)NaCl is an inhibitory substance that can inhibit the growth of certain types of bacteria, and probiotic organisms should tolerate high salt content in the human intestine[24]. It was also reported previously, Lactobacillus species isolated from curd was able to withstand 1-6% of NaCl and excellent growth was recorded in the range 1-5% of NaCl concentration. Isolated lactobacillus species from yogurt samples were also evaluated and their tolerance at different concentrations were recorded [24], [25]. Isolates were successfully grown at (1-9%) NaCl concentrations[26]. This study is also aligned with above-mentioned studies. According to the guidelines of the World Health Organization (WHO) and the Food and Agriculture Organization (FAO, 2002), probiotic LAB need to possess the characteristics of gastric acid and bile resistance as its precondition for probiotics selection. LAB with probiotic potential should have several qualities, such as acid tolerance in the gastrointestinal tract and maintaining a viable physiological state at the site of action [27]. This study reveals isolated probiotic LAB from bovine milk showed similar characteristics/criteria identified by the standard WHO and FAO. Isolates maintained their viability after 24 hours of incubation in acidic condition. These results suggest that isolates have a probiotic property that can live in the GIT can probably survive the passage through the liver and intestines. Tolerance to Bile salt, pancreatic fluid, and stomach pH have an essential role in assuring the viability and development of potential probiotic strain in the gastrointestinal tract[28]. Gastric juice resistance (pH 2-2.5) is a primary functional prerequisite for probiotics that enable selected strains viable through the gastrointestinal tract [29]. The secretion of gastric acid and passage through the stomach is a primary defense mechanism that must be overcome by all ingested microorganisms, including probiotics [30].

Similar findings reported by earlier studies [32–37]. This study confirms previous findings that the acid sensitivity of the LAB does not rely on the source of the sample. For example, the viability percentage of BCM1 (89.9%) from cow’s milk is less than BBM3 (92.7%) buffalo’s milk, whereas BCM2 (94.9%) of cow’s milk has better survival percentage than BBM1 (88.4%) buffalo milk samples (Figure 1 and Table 3). All isolates exhibited tolerance against acidic pH with maximum figures in BCM2 and minimum in BCM3 (Figure 1 and Table 3). Hence this study concludes LAB isolates from bovine milk exhibited tolerance against acidic pH and sodium chloride concentrations. The ability of LAB isolates to withstand acid and sodium chloride salt makes them a potential probiotic bacterium coping up the gastric juice in the GI tract.

The bile tolerance activity result suggests that isolates showed a probiotic property that can live in the GIT can probably survive the passage through the liver and intestines. The potential to withstand bile salt at a concentration of 0.3 percent has physiological importance as it is naturally found in the human intestine [37]. All seven Lactobacillus isolates showed good tolerance of bile at 0.3% of bile salt concentration. This study is in agreement with the previous studies [31], [38]. LAB requires bile tolerance properties in small intestine, where it plays an essential role in specific and non-specific defense mechanisms in order to survive and be functionally successful. The concentration of bile salts is the defining factor of the magnitude of its inhibitory effects which is a necessity for the colonization and metabolism of probiotic bacteria in the host’s small intestine. Concentration of bile salts varies with the time of digestion; 0.3% (w/v) is the average concentration of bile salts in the human bowel and the concentration used to measure LAB probiotic property studies [39]. Generally, LAB from dairy source is probably the best candidates to enhance the nutritional value of foods as probiotic bacteria.

Since significant amounts of deconjugated bile salts may have unwanted side effects on the human host, there may be questions about the safety of a bile salt hydrolase (BSH)-positive probiotic strain. Bacterial genera most suitable to be used as probiotics (bifidobacteria and lactobacilli) are, however, unable to dehydroxylate deconjugated bile salts. Numerous roles of BSH positive LAB including, their nutritional benefit, membrane alterations, bile detoxification, have been reported in recent studies. Bile tolerance and BSH activity of LAB isolates are important to their survival in GIT[11], [40]. This study found a correlation between bile salt tolerance and BSH activity. LAB isolates in the study were found both bile salt-tolerant and BSH positive (Table 4). BSH enzyme role in bile resistance is not yet fully revealed, but it is proposed that, since the non-dissociated form of bile salts that exhibit toxicity by raising intracellular pH as similar fashion as organic acids. The BSH-positive cells may survive by forming the weaker un-conjugated form [11].

Previous studies confirm BSH activity is a common trait in LAB species like *L. plantarum* [41]. Microorganism’s BSH activity has been reported as key factor determining the probiotic ability of strains [42]. Cholesterol-lowering has an association with bile salt hydrolysis [21]. Studies indicate that Lactobacillus’s cholesterol-lowering property is attributed to cholesterol interaction into the cell membranes of bacteria during its growth period. Nonetheless, the main mechanism is related to bile salt hydrolase. It has been proposed that BSHs facilitate the consumption of cholesterol or bile into bacterial membranes [43], [44]. BSH positive strains secrete bile salt hydrolase enzyme that hydrolyzes conjugated bile acids to give deconjugated bile acids and amino acids [31]. The prevalence of BSH activity is primarily a significant factor for GIT occupants of the *Lactobacillus and Bifidobacterium* [45]. In many cases, Lactobacillus from dairy isolates were found to produce functional *in-vitro* BSH enzymes [46]. Generally, BSH activity has an important functional role in a probiotic LAB due to lowering the cholesterol level, strengthening cell membrane due to incorporation of cholesterol to membrane and providing a nutritional role by providing carbon, nitrogen and energy source due to de-conjugation of bile salt[47].

One of the basic criteria to select probiotics is their antimicrobial properties due to the production of compounds such as organic acids, short-chain fatty acids, and bacteriocins. The mechanism of antagonistic activity of the LAB against pathogenic strains is well documented by Sui-skovi’c et al[48]. The ability to generate numerous antimicrobial compounds is one of the main features which enable the probiotic strain to stay competitive and exclude pathogens in the intestine and provide a probiotic benefit for the host. LAB & Bacillus sp. Antimicrobial activity against microorganisms is attributed to various kinds of inhibitory metabolites, mainly bacteriocin[49] [50]. H_2_O_2_ is one of the primary metabolites responsible for antagonistic property of lactic acid bacteria[31]. The production of H_2_O_2_ is also considered desirable criteria for food preservation and serves a gap to the proliferation of pathogenic microorganisms[51]. In this report, LAB isolates were shown to have high inhibitory activity against S.aerues. This finding is consistent with[52][53], reported that antimicrobial compounds developed by Lactobacillus have a great potential to hinder pathogenic microorganisms. As [54]states the bactericidal characters of lactobacilli is mainly due to organic acid production which causes reduction in PH. The acidic conditions in the stomach can even increase this antimicrobial activity. In fact, these probiotic properties may be based on the production of different lactic acid concentrations in the gastrointestinal tract, which hinder the proliferation of Gram-negative pathogens in association with detergents such as bile salts [11]. The antibacterial activity of organic acids is attributed to both the reduction of pH and the undissociated form of molecules. Low external pH has been documented to induce acidic cytoplasm, while non-dissociated lipophilic acid is passively distributed across the membrane. Undissociated acid function by disrupting the permeability of the cell membrane, resulting in an interference in the transport of substrates. Organic acids have a significant inhibitory effect on gram-negative bacteria[55]. Healthy oral microbiota is usually based on ability of probiotics to inhibit the development of oral cavity-related pathogens[56].this study is in support of [57].

## 5.0 Conclusion

The LAB isolates from bovine milk exhibited acid and bile tolerance activity. Since the bacteria must be able to survive acidic conditions in the stomach and avoid bile acids at the beginning of the small intestine to serve as a probiotic to benefit the host, these isolates have probiotic candidates for further *in-vivo* studies. It needs to assess their technological characteristics for application as novel probiotic starters. The findings of this study generally suggest that LAB isolated from bovine milk were resistant to acid, bile and sodium chloride. Future studies on the probiotic ability and antibacterial activity of LAB from bovine milk will make a new breakthrough on the nutritional benefit of bovine milk and LAB. The results of this investigation can also provide baseline information for future studies about the probiotic potential of LAB from bovine milk which could be formulated as a starter culture in food products and further human consumption as a probiotic supplement.

## Acknowledgments

Authors are thankful to Sharda University for providing laboratory facilities along with financial and academic assistance for conducting this research. The support of Wollo University, Ethiopia is also kindly acknowledged.

