## supplemental tables for "Anti-microbial activity of potential probiotic lactic acid bacteria against Methicillin-Resistant Staphylococcus Aureus (MRSA)"

**Table 1A: Morphological, physiological and biochemical characterization of LAB isolates.**

**+=growth - =no growth**

| **Isolates** | **Shape** | **Catalase**  **test** | **Growth at different temperature (°C)** | | | **Gram test result** | **Motility test** | **Surface** | **Margin** | **Elevation** | **Color** |
| --- | --- | --- | --- | --- | --- | --- | --- | --- | --- | --- | --- |
|  |  |  | **15** | **30** | **45** |  |  |  |  |  |  |
| **BCM1** | Cocci | - | + | + | + | + | Non motile | Smooth | Entire | Convex | white |
| **BCM2** | Rod | - | + | + | + | + | Non motile | Rough | Entire | Convex | white |
| **BCM3** | Cocci | - | + | + | + | + | Non motile | Rough | Entire | Convex | white |
| **BBM1** | Tetrad | - | + | + | + | + | Non motile | Smooth | Entire | Convex | white |
| **BBM2** | Rod | - | + | + | + | + | Non motile | Smooth | Entire | Convex | white |
| **BBM3** | Cocci | - | + | + | + | + | Non motile | Smooth | Entire | Convex | white |
| **BGM1** | Cocci | - | + | + | + | + | Non motile | Smooth | Entire | Convex | white |

**Table 1 B: Carbohydrate fermentation result**.

| **Isolates** | **Carbohydrate fermentation** | | | | |
| --- | --- | --- | --- | --- | --- |
|  | **Sucrose** | **Lactose** | **Mannitol** | **Dextrose** | **Maltose** |
| **BCM1** | **+** | + | + | + | + |
| **BCM2** | **+** | + | + | + | + |
| **BCM3** | **+** | + | + | + | + |
| **BBM1** | **+** | + | + | + | + |
| **BBM2** | **+** | + | + | + | + |
| **BBM3** | **+** | + | + | + | + |
| **BGM1** | **+** | + | + | + | + |

**Table 2: Viability Status of LAB isolates at different NaCl concentrations**

**++= high growth + =growth- =no growth**

| **Isolates** | **Viability of LAB isolates at different salt concentrations** | | |
| --- | --- | --- | --- |
|  | **5%NaCl** | **10% NaCl** | **15% NaCl** |
| **BCM1** | + | ++ | - |
| **BCM2** | ++ | ++ | - |
| **BCM3** | ++ | + | - |
| **BBM1** | + | _ | - |
| **BBM2** | + | ++ | - |
| **BBM3** | + | + | - |
| **BGM1** | + | ++ | - |

**Table 3: Acid tolerance of LAB isolates (pH=3) compared to pH=7. Data expressed as mean ± SD (n = 3).**

| **Lactobacillus species** | **Cell viability (log CFU/ml)^1^** | |  |
| --- | --- | --- | --- |
|  | **pH 7.2** | **pH 3.0** | **Cell viability percentage (%)** |
| **BCM1** | 7.47±0.07 | 6.72±0.21 | 89.9 |
| **BCM2** | 7.28±0.09 | 6.91±+0.12 | 94.9 |
| **BCM3** | 7.72±0.18 | 6.75±0.12 | 87.4 |
| **BBM1** | 7.90±0.04 | 6.99±0.03 | 88.4 |
| **BBM2** | 7.54±+0.05 | 6.65±0.18 | 88.1 |
| **BBM3** | 7.54±+0.04 | 6.99±0.03 | 92.7 |
| **BGM1** | 7.48±0.10 | 6.87±0.07 | 91.8 |

**Table 4: Bile salt hydrolase activity of LAB isolates**

**+ positive**

| **Isolates** | **Bile Salt Hydrolase activity** |
| --- | --- |
| **BCM1** | + |
| **BCM2** | + |
| **BCM3** | **+** |
| **BBM1** | + |
| **BBM2** | + |
| **BBM3** | + |
| **BGM1** | + |
